# Fish Aggregating Devices could enhance the effectiveness of blue water MPAs

**DOI:** 10.1101/2022.11.16.516764

**Authors:** Michael Bode, Edward T Game, Alex Wegmann, Kydd Pollock

## Abstract

In the past two decades, drifting fish aggregation devices (FADs) have revolutionised pelagic fisheries, and are now responsible for the majority of tuna purse seine catches. Here, we argue that by taking advantage of the same proven aggregative properties, FADs could be used to enhance the benefits provided by blue water Marine Protected Areas (MPAs). Using models of commercially-targeted fish populations, we explore the potential benefits that could be achieved if unfished conservation FADs were positioned within blue water MPAs. Our results suggest that conservation FADs could deliver benefits, both to target species and the broader ecosystem. By increasing the residence time of exploited species, conservation FADs will reduce average mortality rates inside MPAs. By increasing the local density of species whose populations are depressed by exploitation, FADs can also improve the function of ecosystems in blue water MPAs. Conservation FADs could therefore amplify the benefits of blue water MPAs. We find this amplification is largest in those contexts where blue water MPAs have attracted the most criticism - when their area is small compared to both the open ocean and the distribution of fish stocks that move through them.

## INTRODUCTION

The open-ocean is under unprecedented threat from human activities. The creation of blue water marine protected areas (MPAs; Figure 1) has been an important part of the response to this threat (Wagner 2013). Blue water MPAs are large-scale (>100,000 km2) protected areas that encompass open-ocean, pelagic ecosystems, although they are often centred upon oceanic islands, reefs, or seamounts. The number and extent of blue water MPAs has accelerated rapidly in recent years and is likely to continue to accelerate in support of international agreements such as the Convention on Biological Diversity’s post-2020 Global Biodiversity Framework, and the ongoing negotiations around the conservation and sustainable use of Biodiversity Beyond National Jurisdiction.

**Figure 1:**
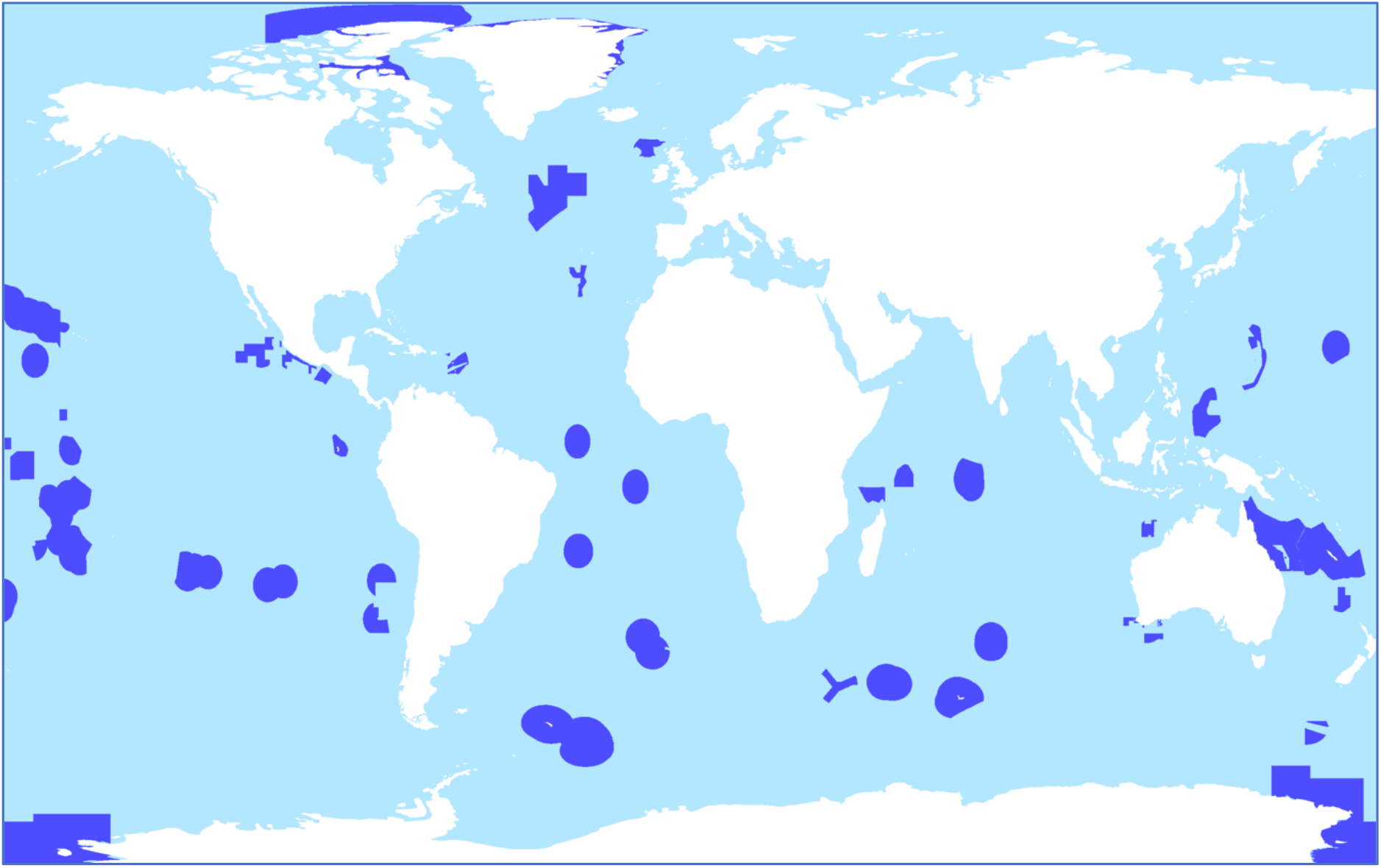
Distribution of the world’s blue water MPAs (shown in dark blue), defined as marine reserves with areas larger than 100,000 square kilometres. Data on MPA boundaries are sourced from the World Database of Protected Areas (UNEP-WCMC 2014).

The anticipated ecological benefits of blue water MPAs are uncertain for two reasons. First, despite being among the world’s largest protected areas, blue water MPAs still only cover a tiny fraction (<5%) of the ocean’s vast area (O’Leary et al. 2018). Their ecological benefits may be commensurately small. Second, a substantial proportion of pelagic biodiversity is highly mobile or migratory and can easily move across reserve boundaries. This is especially true for species targeted by commercial fisheries: including tuna and swordfish (Boerder et al. 2019), whose migration patterns may encompass whole oceans. Even the largest blue water MPAs could not protect individuals of these species from fishing mortality throughout their lives, a key design objective of coastal and reef MPAs (Green et al. 2015). As a consequence, the ability of blue water MPAs to conserve pelagic species and ecosystems is a question of intense debate (Game et al. 2009; Kaplan et al. 2010; Koldewey et al. 2010; Gilman et al. 2019, 2020).

Drifting Fish Aggregation Devices (dFADs) are floating objects deployed by fishers, to exploit the natural propensity of many pelagic species to congregate under such objects. By increasing the local density and consistency of target species, dFADs increase fishers’ expected catch-per-unit-effort (Hanich et al. 2019). Although dFADs attract a variety of pelagic species, their effectiveness at aggregating commercially important tropical tuna species has made them the mainstay of the world’s purse seine tuna fisheries (Maufroy et al. 2017). Drifting FADs increase local pelagic biomass density, and are increasingly fitted with echosounder-equipped, satellite-linked buoys. These provide real-time information on the biomass around them, greatly increasing the efficiency of fisheries production (Baidai et al. 2020; Wain et al. 2021). The number of active dFADs deployed in the world’s oceans is notoriously difficult to estimate, but likely exceeds 100,000 per year, and is increasing (Gershman et al. 2015). A number of possible negative ecological impacts have been suggested as a result of extensive dFAD use, starting with unsustainable catch levels for target species, but also including bycatch of aggregating species such as silky sharks (Clavareau et al. 2020), detrimental effects on tuna behaviour (Dagorn et al. 2013), and damage caused by abandoned dFADs colliding with vulnerable habitat such as coral reefs (Imzilen et al. 2021). Drifting FADs may also pose a challenge for the effectiveness of blue water MPAs because they cross into and out of the protected areas, taking apex pelagic species with them (Curnick et al. 2021a).

Fish aggregating devices have proven uses in marine conservation. For more than 20 years, anchored FADs have been deployed offshore of fishing communities by development and fisheries organizations, to redistribute fishing pressure from coastal ecosystems to pelagic resources (Campbell et al. 2016; Jauharee et al. 2021). Here, we propose deploying FADs that remain inside blue water MPAs could directly and positively enhance their benefits. For the purpose of this paper, we refer to these unfished FADs remaining inside blue water MPAs as conservation FADs (cFADs). These cFADs differ from anchored FADs in in fundamental ways: first, they are not fished. Second, they do not need be anchored to the seafloor to remain inside blue water MPAs, rather we imagine them as self-powered devices capable of remaining semi-stationary.

cFADs could deliver benefits to species that experience direct fishing mortality, and also to the broader ecosystem. By increasing the residence time of highly-mobile and migratory species inside the protected area, cFADs decrease the exposure of these species to fishing mortality. Despite the temporary nature of this protection, it could nevertheless enhance stock levels, via the same mechanism as temporary closures. An increased residence time could deliver density-dependent ecosystem benefits, which are lost when commercial fishing reduces the abundance of target or bycatch species. For instance, bird species often rely on high tuna densities for effective foraging (Au & Pitman 1986; Jaquemet et al. 2005).

The deployment of cFADs inside blue water MPAs could therefore help to counterbalance the extensive use of dFADs by fisheries in the open ocean, allowing MPAs to enhance their benefit without increasing their size. However, as far as we are aware, the potential for cFADs to amplify the benefits of blue water MPAs has not previously been explored, theoretically or empirically. The goal of this study is to explore the logic of this proposal, by introducing cFADs to standard theoretical models of exploited fish species with MPAs.

## NONSPATIAL MODEL

### Nonspatial model

The first model describes the effects of cFADs as simple, spatially-implicit redistributions of stock numbers. Following ideas from optimal foraging theory (Visser & Fiksen 2013), we assume that each FAD attracts fish by offering them greater benefits than the surrounding seascape. As a consequence, all FADs simply concentrate existing animals from the surrounding seascape. This aggregative role is independent of their spatial location – they aggregate fish both inside and outside protected areas – and this means that dFADs could play a positive conservation role when they drift inside the MPA.

A stock of fish has population *X*, which is distributed across an area *N*, of which an area *M* has been placed into a blue water MPA. The area contains fishing dFADs, which move at random across the entire stock distribution (including some time spent inside the MPA). The area also contains *f_c_* stationary cFADs, all of which are permanently inside the MPA.

Fish gain some density-dependent benefit from being in proximity to any FAD, and the population therefore aggregates in increasing density at each FAD until the marginal benefit of joining a FAD aggregation falls to zero (24). We use the parameter *x* « *X* to denote the proportion of the total fish stock that is found in the vicinity of each FAD. As *x* increases, the aggregating effect of the FADs increases, meaning a higher density of fish will be found in the vicinity of each device. Those fish that are not associated with any FAD will be distributed evenly across the stock distribution area *N*.

In the absence of any FADs, the time-averaged number of fish that will be found inside the MPA will then be:

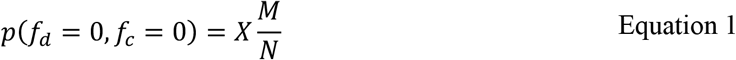

Equation 1 represents the essential role of an MPA – to protect a proportion of the fish stock distribution from mortality. It is also the basis of the primary critique of blue water MPAs: that their size is small relative to the area of a mobile pelagic stock (i.e., *M* ≪ *N*). If cFADs and dFADs are present, then the time-averaged proportion of the stock that will be found inside the MPAs (and therefore protected from excess fishing mortality) is:

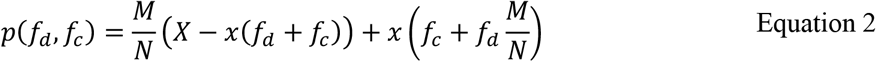

The first term on the right-hand side of Equation 2 represents fish that are not associated with any FAD and are protected by the MPA, while the second term represents those fish that are associated with any FAD, and which are currently inside the MPA. We also assume that *x* ≤ *X*/(*f_d_* + *f_c_*), to ensure non-negative fish populations.

Thus, the presence of *f_c_* conservation cFADs amplify the stock benefits flowing from an MPA by a factor *A*:

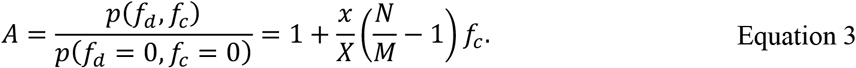

For example, an amplification factor of *A* = 1.5 indicates that the addition of cFADs increases the benefits of the MPA by 50%, compared to the MPA without any cFADs.

We parameterised the nonspatial model with area *N* = 1 and a normalised stock abundance of *X* = 1. We allow the MPA to represent 2%, 4%, and 8% of the stock distribution, ranging between the current global coverage of strong to fully-protected marine reserves (2%), and the current global coverage of all proposed and implemented marine reserves (8%) (Sala et al. 2018). We allow the number of cFADs to range between 0 ≤ *f_c_* ≤ 10. A proportion *x/X* = 10^-3^ (i.e., 0.1%) of the total stock abundance are found in the vicinity of each FAD (of either type). Note that the amplification factor *A* is not influenced by *f_d_*, the number of dFADs.

### Nonspatial model results

Figure 2 shows that the amplification factor *A* is an inverse function of MPA size – highest for smaller MPAs (Figure 2A). Small MPAs offer the least protection to fish stocks (Equation 1), and their smaller benefits are therefore easy to amplify. If a particular number of cFADs create an amplification factor of *A* = 2, for example, then they have effectively doubled the density of targeted fish, halved the exposure of each fish to excess fishing mortality, and doubled the effective size of the MPA, when compared to an MPA without any cFADs present. Figure 3 provides a visual illustration of how cFADs amplify the effective size of a blue water MPA.

**Figure 2:**
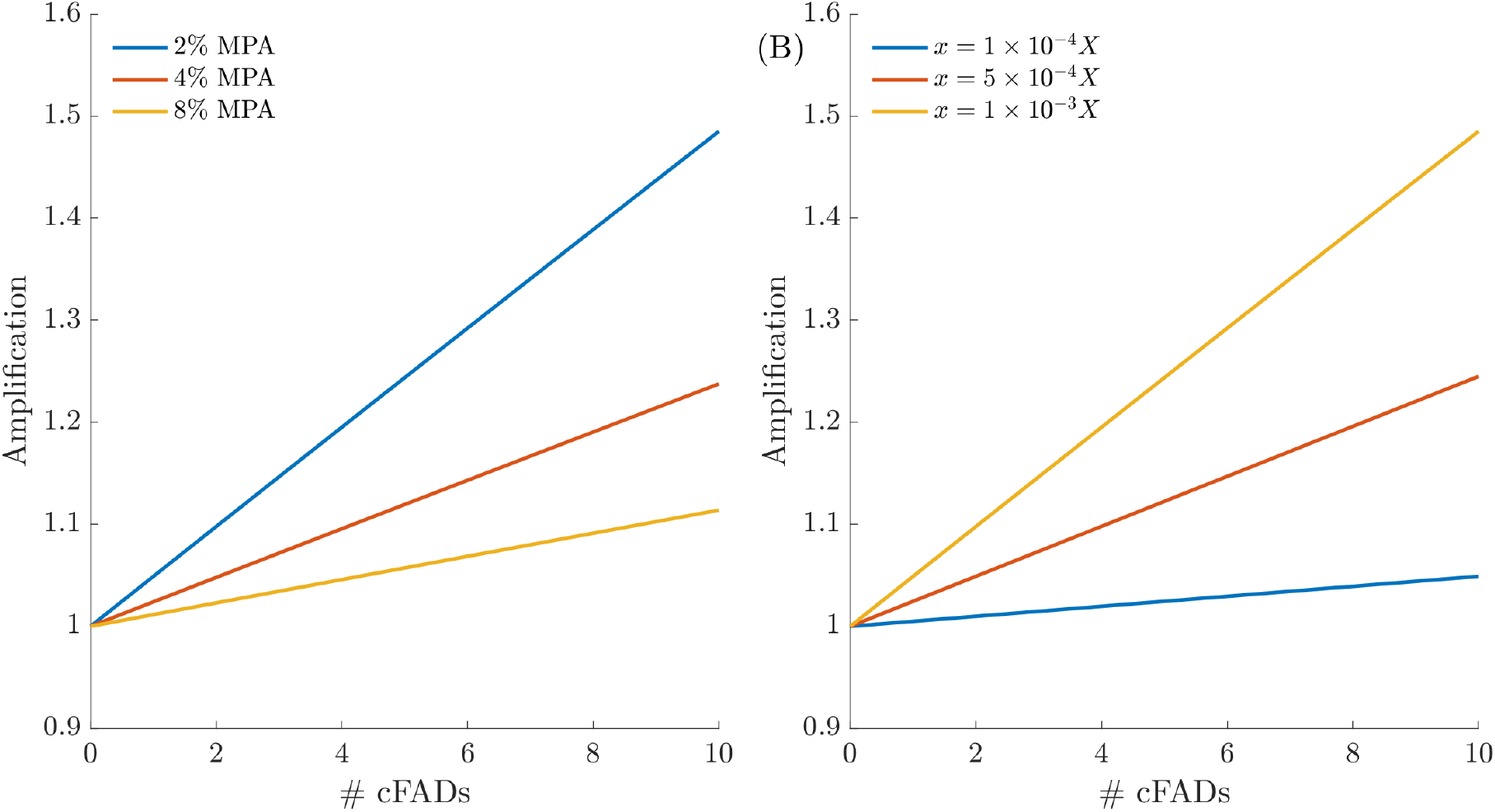
Nonspatial model results. Panel (A) shows the amplification provided to blue water MPAs of three different sizes, as the number of cFADs increases from zero to 10 (x = 1 × 10^-3^ in this panel). Panel (B) shows the amplification provided when FADs attract more fish, as the number of cFADs increases from zero to 10 (with a blue water MPA that encompasses 2% of the total area).

**Figure 3:**
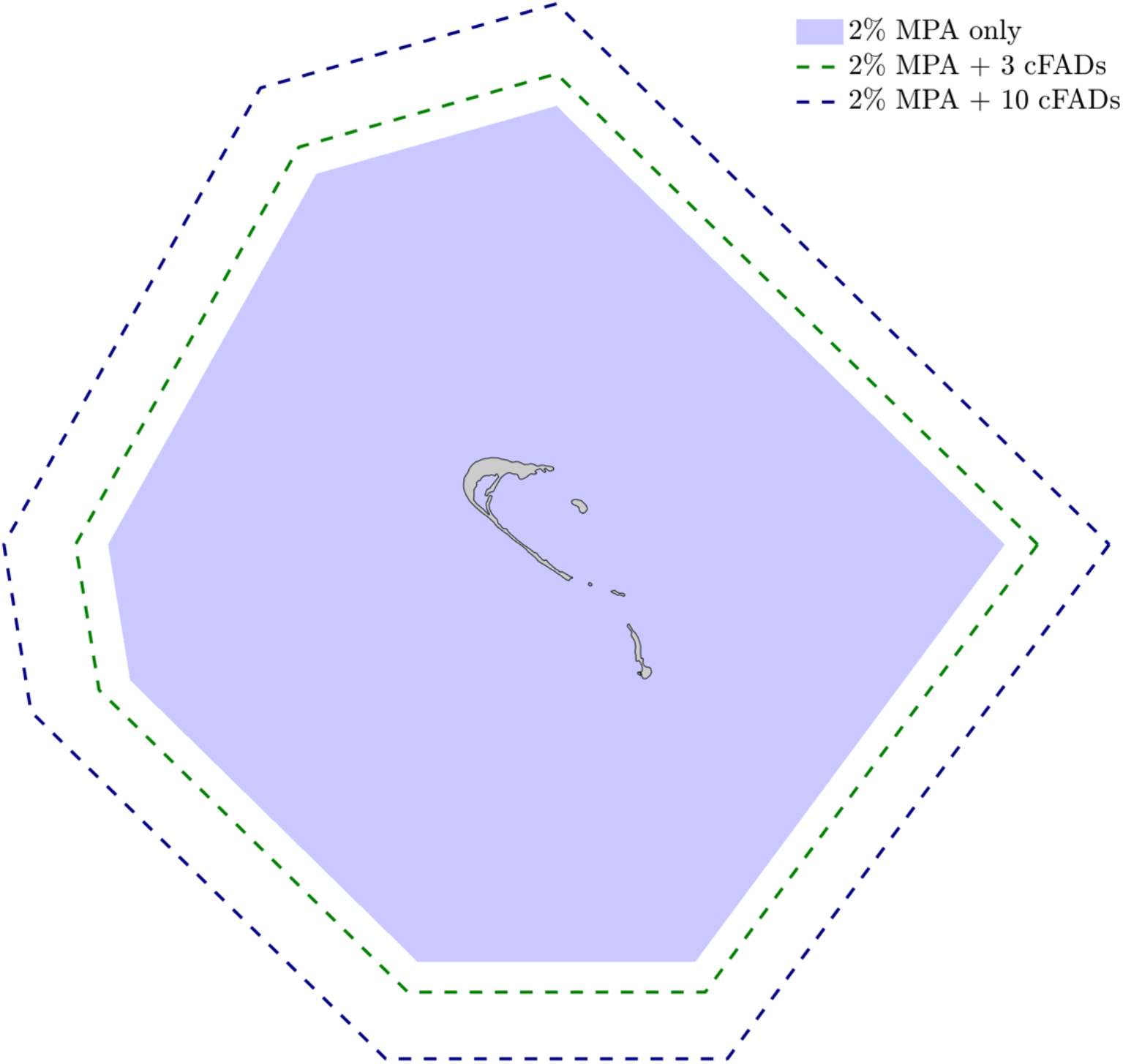
Visual illustration of the benefits provided by cFADs to a hypothetical blue water MPA around a coral atoll. The blue region shows a hypothetical blue water MPA surrounding an island (grey outline). The island us for illustrative purposes only; it has no influence on the model results. With 3 cFADs, inside a 2% MPA, the area will contain as many fish as an MPA that is 15% larger (area enclosed by green dashed line). With 10 cFADs, the MPA will contain as many fish as an MPA that is 50% larger (area enclosed by blue dashed line). The stock distribution area is much larger than domain shown here. Results are based on our nonspatial model in Figure 2A.

Since dFADs compete to attract the same fish as cFADs, our intuition suggested that more dFADs deployed for fishing would result in a net export of fish from the MPA. We therefore expected that a larger number of dFADs would reduce the amplification. However, this turned out not to be the case – the solution for *A* (Equation 3) did not contain any *f_d_* terms. This is because dFADs spend time inside the MPA as well as outside the MPA (e.g., they would spend 10% of their time in an MPA that covered 10% of the stock area), and they therefore do not affect the distribution of the fish stock across the MPA boundary.

While this model captures the attraction that FADs exert on fish population distributions, it doesn’t explicitly model the interaction of FADs with MPA locations (particularly the potential for dFADs to export biomass from no-take zones as they pass through them or along boundaries), nor does it include the negative effects of fishing mortality on stock abundance, which will increase with the number of fishing FADs. To address these factors, we extend the model to include the effects of space, and the negative demographic effects of fishing.

## SPATIAL MODEL

### Spatial model formulation

Our spatial model is based on a previously-published model designed to measure the effects of FADs on the movement of skipjack tuna (Kleiber & Hampton 1994). We assume a one-dimensional seascape that is divided into *n* discrete cells, mapped onto a circle to avoid boundary conditions. A contiguous subset of *m* cells in the centre of the domain are designated as a blue water no-take MPA. We assume that in *f_d_* of the cells across the domain there is a drifting dFAD, and in *f_c_* of the cells inside the MPA there is a stationary cFAD.

Within the domain there is a population of fish that is expected to benefit from the cFADs and MPA. The abundance of these fish in cell *i* at time *t* is denoted *x_i,t_*. Individuals in any cell can move left or right, and this may involve entering or exiting the MPA. They can also remain in the same cell. The relative attractiveness of each cell is defined as *a^i^*, and we assume all cells containing FADs (conservation or fishing) have a higher attractiveness, cells without FADs have a uniform, lower attractiveness, and cells containing multiple FADs have the same attractiveness as single-FAD cells. The result is an accumulation of fish in cells occupied by a cFAD or a dFAD, and a minor accumulation of fish in nearby cells.

Individuals face a constant natural mortality probability of *μ*, and we apply a constant excess fishing mortality rate *e_i,t_* to all cells outside the MPA that contain dFADs in timestep *t*. The mortality rate increases proportionally when multiple dFADs co-occur in space. The number of new recruits each year is determined by a density-dependent Ricker model with rate parameter *k*, following fishing mortality, natural mortality, and movement. Recruitment from the larval pool is equally redistributed across the domain, with each cell receiving *r_t_* recruits in timestep *t*, where:

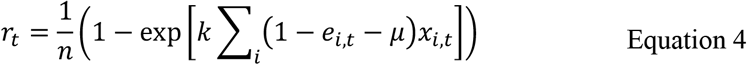

We model reproductive output as a function of local densities rather than global abundance, since density-dependent processes may operate either locally or globally, but this choice does not substantially affect our outcomes. Taken together, these assumptions imply that the abundance of fish in cell *i*. at timestep *t* + 1 is:

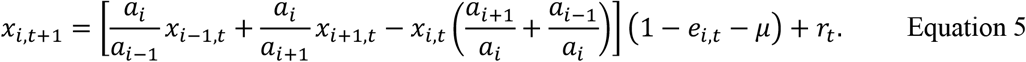

The system does not reach a spatiotemporal equilibrium because the movement of the dFADs constantly affects the distribution of fishes. We therefore simulate the model until the 100-year time-averaged abundance of the total fish population stabilises.

For our example, we discretise the seascape into *n* = 300 sites, with an MPA that takes up *m* = {2,3,6} contiguous sites. We model *f_d_* = {25,150,500} moving dFADs, and *f_c_* = {0,…,10} stationary cFADs. The dFADs are initialised at random outside the MPA, and the cFADs are placed at random locations within the MPA. At the higher densities, many locations will therefore contain more than one FAD. The relative attractiveness of each location is *a^i^* = 0.5 for locations without FADs, and *a^i^* = 0.6 for locations with FADs. The attractiveness of dFADs would seem an important parameter value; however the qualitative form of our results are robust to this choice, as shown in *Supplementary Figure S1*. Natural mortality is assumed to be *μ* = 0.05, fishing mortality is *e_i_* = 0.1, and density-dependent Ricker mortality is governed by parameter *k* = 0.1. The model assumes that fishing mortality in cells without a dFAD is zero, but we also investigated the effects of non-dFAD associated mortality outside the MPA. The model also assumes that fishing mortality on dFADs is constant. Sensitivity analyses show that neither choice affects our results (*Supplementary Figure S2 & S3*).

### Spatial model results

The spatial model results augment the nonspatial results in three important ways. First, unlike the nonspatial model, spatial amplification benefits are concave; that is, increasing the number of cFADs delivers diminishing marginal conservation returns. In the nonspatial model, each cFAD delivers independent benefits, drawing fish into the MPA from across the fished area. In the spatial model by contrast, cFADs can only attract fish from the surrounding seascape. As their numbers increase, these local areas begin to overlap, the cFADs begin to compete for fish density. Such a saturation effect has been observed for skipjack tuna in the Solomon Islands (Kleiber & Hampton 1994).

Second, unlike the nonspatial model, the number of dFADs outside the MPA negatively affects the benefits produced by cFADs inside the MPA (Figure 4B). In the nonspatial model, dFADs generate benefits to MPAs when they are inside their boundaries, and equal disbenefits when they are outside. By contrast, in the spatial model these benefits and disbenefits are unequal. dFADs enter the MPA with a lower number of associated fish and leave with a larger number: a net export. This inequality is greatest for large MPAs, since dFADs will spend more time traversing them, accumulating a higher biomass in the process.

**Figure 4:**
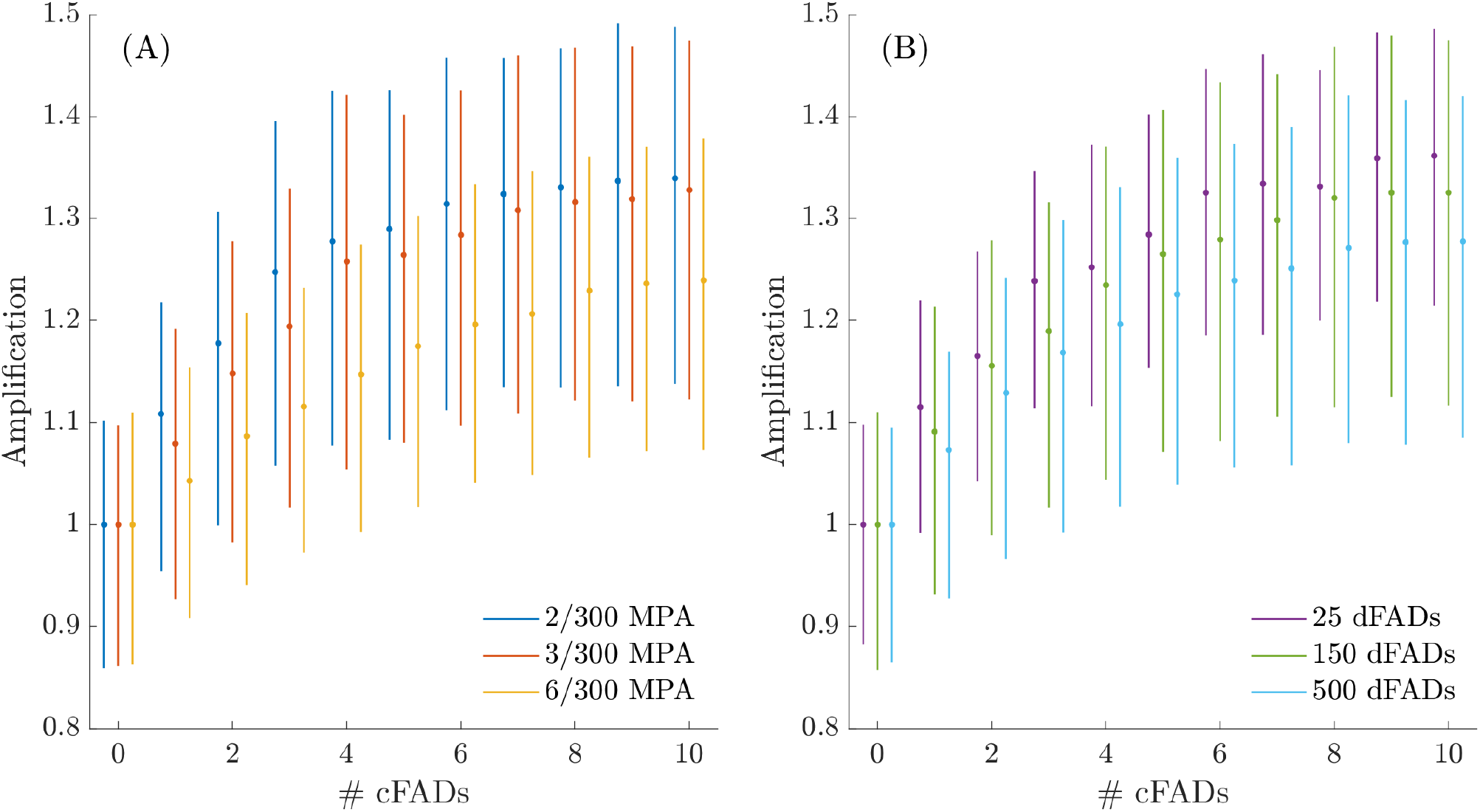
Spatial model results for the number of cFADs varying between 0 and 10. Panel (A) shows that the amplification benefits provided to a blue water MPA decline as the MPA increases in size. There are 150 dFADs in this system. Panel (B) shows that benefits decline as the number of dFADs increases. These results are for an MPA that encompasses 3 cells. In both panels, the bounds illustrate the interquartile range of the observed amplification across the different cells of the MPA, and across many years. That is, how the increase in density inside the MPA varies across space and through time.

Third, the benefits provided by the cFADs vary in space and time, depending on the location of the different FADs and their recent history (e.g., if they have recently exited an MPA). As a consequence, the amplification provided by the cFADs varies in space and time, as captured by the bounds in Figure 4. Although cFADs offer an average amplification, certain locations within the MPA offer higher and lower benefits. For the species and ecosystem functions that benefit from high fish densities (e.g., hunting seabirds), this variation makes it more likely that benefits will be available at some location in the MPA.

## DISCUSSION

The world’s tuna fishing fleets increasingly rely on dFADs to enhance their catches. By producing and maintaining high densities of fish in predictable locations, these fleets use dFADs to increase the efficiency of their catches. Conservation FADs would operate under similar principles, but for a diametrically opposite motivation. By increasing the residence time of species within MPAs, cFADs could increase the efficiency of blue water MPAs, substantially amplify the benefits that they produce. The addition of cFADs to blue water MPAs could address their greatest limitation – that MPAs in the open ocean are small relative to the distribution of targeted stocks. Faced with the goal of increasing the benefits of blue-water MPAs, managers might find it preferable – in terms of financial expense, or perhaps of conflict with fisheries – to add cFADs to an existing MPA, rather than increase its area.

Conservation FADs do not have to reduce the overall mortality risk faced by pelagic fishes to deliver a benefit to blue water MPAs – the ecosystems within an individual blue water MPA can still benefit from an increased local density of aggregating fishes, even if the total abundance of those fishes is the same. Drifting objects, which FADs mimic, help tuna form and maintain school integrity and size (Dagorn et al. 2013), which may increase their feeding efficiency and their visibility to other species that feed in association with them. For example, in the Pacific Remote Islands Marine National Monument, a blue water MPA in the central Pacific, sooty terns feed obligately with yellowfin tuna schools, and great frigatebirds and red-footed boobies benefit from schooling subsurface predators, including tuna species, for effective foraging (Gilman et al. 2019). Foraging efficiency is tightly linked to breeding success, placing a premium on consistent access to resources within the foraging range of the species.

The effectiveness of FADs as fishing gear may be a reason for caution when deploying them for conservation goals. The presence of semi-stationary cFADs in blue water MPAs may make them more attractive to illegal fishers, for example. This risk seems low, however, since illegal fishers would find FADs hard to locate without access to their GPS transponders. Additionally, blue water MPAs are being continually traversed by fishing dFADs (Curnick et al. 2021b), of known location, and the addition of a relatively small number of cFADs will not substantially increase the attractiveness of illegal fishing.

Our results provide a theoretical rationale for deploying cFADs to blue water MPAs. However, the models that we use here are limited in a number of important ways that may affect the potential benefits offered by cFADs. First, cFADs will increase the densities of any species that is attracted to pelagic floating objects, and not all of these will be negatively affected by fishing effort or bycatch mortality. Over 55 bony fish species are frequently found around dFADs used by the tropical tuna purse seine fishery, and many of them will be abundant, fast-growing, and of low conservation concern (Amandé et al. 2008). Second, our models are based on the assumption that the aggregating effects of cFADs and dFADs are equivalent. dFADs move passively with ocean currents, which may place them to current boundary locations that offer better conditions for environmental factors and food. Third, our models are not parameterised for any particular fish species or location; as a consequence, they cannot accurately predict fish densities around cFADs, nor how these densities would change with the number or location of the cFADs. Given the novelty of the cFAD concept, anticipating parameter values for a particular location and species would be very difficult. Our models’ goal was to provide a theoretical justification for the experimental trials that would be needed to estimate these values.

The immediate policy question for conservation FADs in blue water MPAs is whether their benefits are large enough to justify their deployment. Moving forward, we believe that the best method to answer this question is small-scale experimental deployment of cFADs in existing blue water MPAs, with particular target species and ecosystem functions being identified and measured. The downside risks of such experimental management are manageable: MPAs are already being traversed by much larger numbers of dFADs, and monitored cFADs can easily be removed if negative consequences arise. The enthusiasm and speed with which dFADs have been adopted by pelagic fisheries speaks to the size of the potential opportunity.

## ACKNOWLEDGEMENTS

This study was supported by funding from the Anthropocene Institute.

