## Supporting Information for "Fish Aggregating Devices could enhance the effectiveness of blue water MPAs"

**Supporting Figure S1: Spatial model predictions when FADs have different attractiveness.**

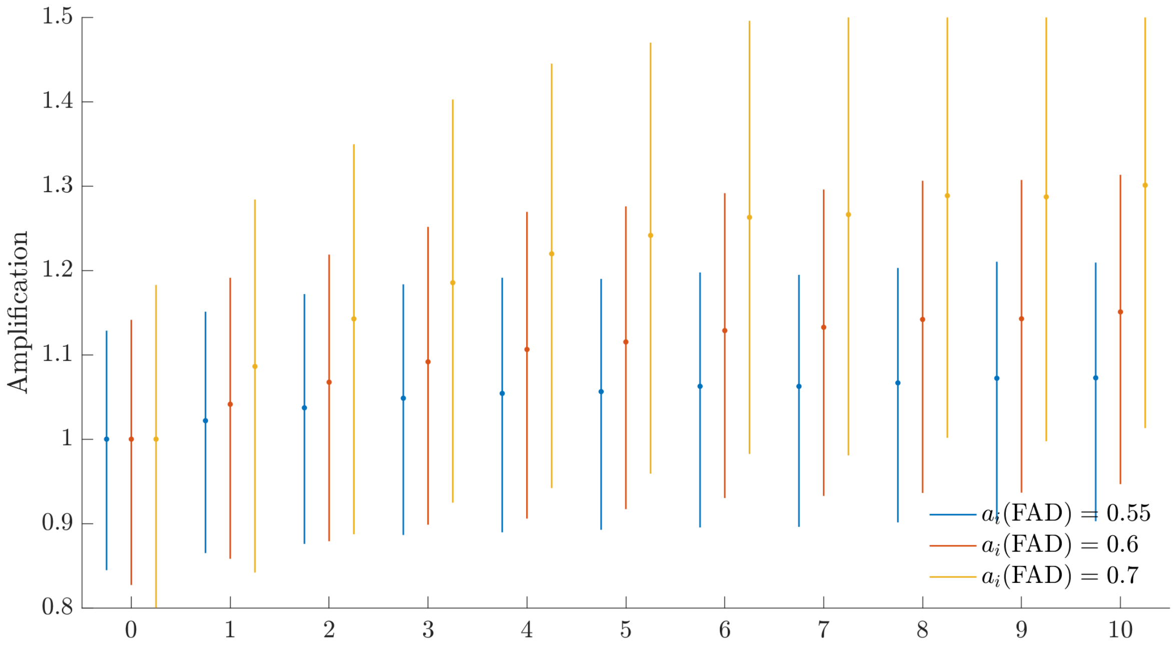


***Supporting Figure S1**: Spatial model results for the number of cFADs varying between 0 and 10. Colours indicate results when FADs have different attractiveness to fish, relative to parts of the ocean without FADs. The same increasing amplification is apparent. Results are for an MPA that encompasses 3 cells (i.e., 1% of the domain). The bounds illustrate the interquartile range of the observed amplification across the different cells of the MPA, and across many years.*

**Supporting Figure S2: Spatial model predictions with fishing mortality in fished areas without dFADs.**

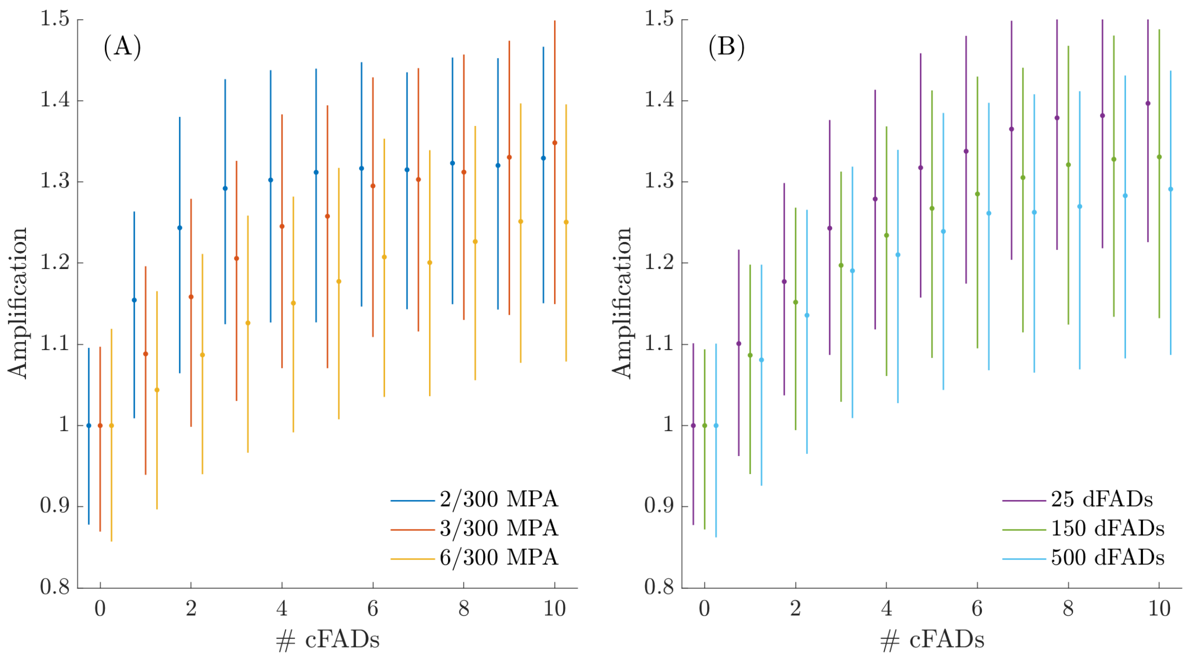


***Supporting Figure S2**: Spatial model results for the number of cFADs varying between 0 and 10. In this figure (as compared to Figure 4 in the main text), the fished population is exposed to fishing mortality in areas without dFADs, as well as in areas with dFADs (assuming in both cases that the area is not inside a blue water MPA). In particular, the background fishing mortality rate is 0.01 per timestep, and the dFAD fishing mortality rate is 0.1 per timestep. The introduction of background mortality to the model has negligible effects on the results.*

*Panel (A) shows that the amplification benefits provided to a blue water MPA decline as the MPA increases in size. There are 150 dFADs in the system. Panel (B) shows that benefits decline as the number of dFADs increases. These results are for an MPA that encompasses 3 cells (i.e., 1% of the domain). In both panels, the bounds illustrate the interquartile range of the observed amplification across the different cells of the MPA, and across many years.*

**Supporting Figure S3: Spatial model predictions with fishing mortality in fished areas without dFADs.**

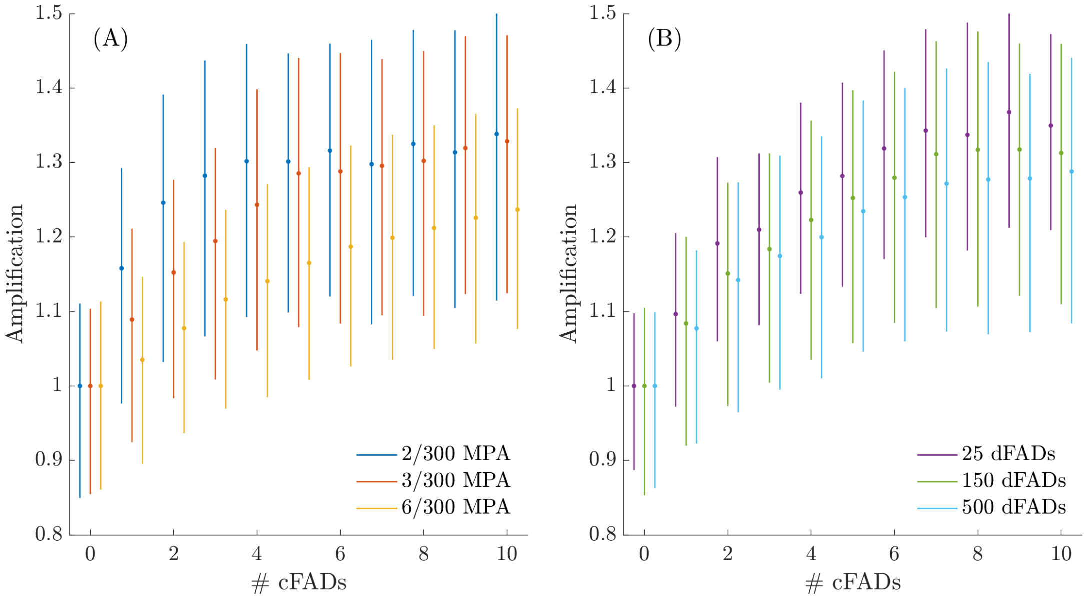


***Supporting Figure S3****: Spatial model results for the number of cFADs varying between 0 and 10. In this figure (as compared to Figure 4 in the main text), areas with dFADs that are not inside a blue water MPA experience a variable dFAD fishing mortality rate. Specifically, the value of* $e_{i,t}$ *in the model is either 0.2 per timestep (with a 50% probability), or is zero. This mimics dFADs which are not harvested every timestep. The difference between the results is negligible.*

*Panel (A) shows that the amplification benefits provided to a blue water MPA decline as the MPA increases in size. There are 150 dFADs in the system. Panel (B) shows that benefits decline as the number of dFADs increases. These results are for an MPA that encompasses 3 cells (i.e., 1% of the domain). In both panels, the bounds illustrate the interquartile range of the observed amplification across the different cells of the MPA, and across many years.*

**Supporting Figure S4: Spatial model predictions with fishing mortality in fished areas without dFADs.**

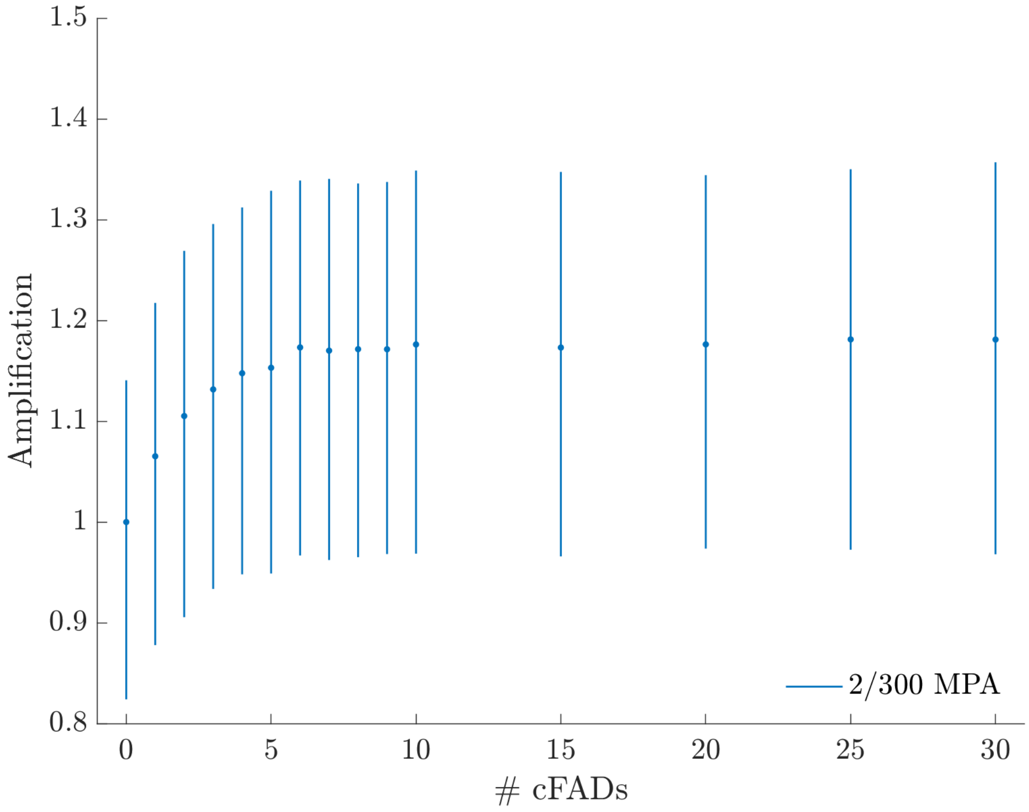


***Supporting Figure S4****: Figure 4: Spatial model results for the number of cFADs varying between 0 and 30, to better show the asymptotic benefits of more cFADs. The amplification benefits provided to a blue water MPA decline as the MPA increases in size. There are 150 dFADs in the system. Bounds illustrate the interquartile range of the observed amplification across the different cells of the MPA, and across many years.*
